# Adaptation to larval crowding in *Drosophila ananassae* and *Drosophila nasuta nasuta*: increased larval competitive ability without increased larval feeding rate

**DOI:** 10.1101/011684

**Authors:** Archana Nagarajan, Sharmila Bharathi Natarajan, Mohan Jayaram, Ananda Thammanna, Sudarshan Chari, Joy Bose, Shreyas V. Jois, Amitabh Joshi

**Author notes:** equal contribution (Archana Nagarajan, Sharmila Bharathi Natarajan).

## Abstract

The standard view of adaptation to larval crowding in fruitflies, built on results from 25 years of multiple experimental evolution studies on *D. melanogaster*, is that enhanced competitive ability evolves primarily through increased larval feeding and foraging rate, at the cost of efficiency of food conversion to biomass, and increased larval tolerance to nitrogenous wastes. These results, moreover, were at odds from the predictions of classical *K*-selection theory, notably the expectation that selection at high density should result in the increase of efficiency of conversion of food to biomass, and were better interpreted through the lens of *α*-selection. We show here that populations of *D. ananassae* and *D. n. nasuta* subjected to extreme larval crowding evolve greater competitive ability and pre-adult survivorship at high density primarily through a combination of reduced larval duration, faster attainment of minimum critical size for pupation, greater time efficiency of food conversion to biomass, increased pupation height with a relatively small role of increased urea/ammonia tolerance, if at all. This is a very different suite of traits than that seen to evolve under similar selection in *D*. *melanogaster* and seems to be closer to the expectations from the canonical theory of *K*-selection. We discuss possible reasons for these differences in results across the three species. Overall, the results reinforce the view that our understanding of the evolution of competitive ability in fruitflies needs to be more nuanced than before, with an appreciation that there may be multiple evolutionary routes through which higher competitive ability can be attained.

## Introduction

Adaptation to crowding leading to greater competitive ability, first conceptualized as *r*-and *K*-selection theory (MacArthur and Wilson 1967), is an important phenomenon in ecology and evolution that was extensively studied in *Drosophila melanogaster* over the past three decades, using experimental evolution approaches (reviewed in Joshi 1997; Mueller 1997; Prasad and Joshi 2003). With lifespan, fecundity, starvation resistance and dessication resistance, some of the other traits extensively examined via experimental evolution in *D. melanogaster*, there were conflicting reports of differing patterns of correlated responses to selection between laboratories (Ackermann et al. 2001; Prasad and Joshi 2003; Rose et al. 1996), that led, in part, to a growing appreciation that evolution is often “local” (Rose et al. 2005). In the case of adaptation to crowding, however, the pattern of correlated responses to selection was largely consistent over two separate selection experiments, involving *D. melanogaster* populations originating from different geographical sources (*r*- and *K*-populations: Mueller and Ayala 1981; UU and CU populations: Joshi and Mueller 1996), and the suite of traits through which selected populations evolved greater competitive ability was very different from the canonical version of *r*- and *K*-selection theory that emphasized the centrality of increased food to biomass conversion efficiency as an adaptation to chronic crowding (Mueller 2009).

Results from the earlier selection studies on adaptation to crowding in *D. melanogaster* have been reviewed extensively (Joshi 1997; Mueller 1997; Joshi et al. 2001; Prasad and Joshi 2003; Dey et al. 2012) and, consequently, we will just summarize the major observed correlated responses of pre-adult traits to selection. Relative to the low density *r*-populations, the crowding adapted *K*-populations exhibited greater (i) pre-adult survivorship, when assayed at high larval density (Bierbaum et al. 1989), (ii) pre-adult competitive ability (Mueller 1988), (iii) larval feeding rate (Joshi and Mueller 1988), (iv) larval foraging path length (Sokolowski et al. 1997), (v) pupation height (Mueller and Sweet 1986; Joshi and Mueller 1993), and (vi) minimum food requirement for pupation (Mueller 1990). Pre-adult survivorship of the *r*- and *K*-populations did not differ significantly when assayed at low larval density (Bierbaum et al. 1989). Pre-adult development time in the *K*-populations was lower than that of the *r*-populations at low (30 larvae per vial) and moderately high (160 larvae per vial) density, but greater at very high (320 larvae per vial) density (Bierbaum et al. 1989). The evolution of higher larval feeding rate and pupation height in crowding adapted populations was subsequently verified with a separate set of populations (*rK* and *r*×*rK* populations), by Guo et al (1991).

Relative to the control UU populations, the larval crowding adapted CU populations exhibited greater (i) pre-adult survivorship, when assayed at high larval density (Mueller et al. 1993; Shiotsugu et al. 1997), (ii) larval feeding rate (Joshi and Mueller 1996), (iii) larval foraging path length (Sokolowski et al. 1997), (iv) minimum food requirement for pupation (Joshi and Mueller 1996), and (v) tolerance to nitrogenous wastes like urea (Shiotsugu et al. 1997; Borash et al. 1998) and ammonia (Borash et al. 1998). Pre-adult survivorship of the CU and UU populations did not differ significantly when assayed at low larval density (Mueller et al. 1993; Shiotsugu et al. 1997). CU pre-adult development time was similar to the UU populations when assayed at low larval density, but lower than the UU populations when assayed at high larval density (Borash and Ho 2001). Pupation height of the CU populations was greater than the UU controls during early generations of CU selection (Mueller et al. 1993), but did not differ significantly from the controls after about 60 generations of selection (Joshi and Mueller 1996); possible reasons for this are extensively discussed by Joshi et al. (2003). Urea and ammonia tolerance were not assayed on the *r*- and *K*-populations, whereas competitive ability was not assayed on the CU and UU populations.

The *K*-populations differed from the *r*-populations in experiencing higher larval as well as adult density and, moreover, were kept on an overlapping generation regime whereas the *r*-populations were kept on discrete generations (Mueller and Ayala 1981). The CU populations, on the other hand, differed from the UU controls only in larval density (Joshi and Mueller 1996). Given the congruence in the pre-adult traits that evolved in the CU and *K*-populations, it is likely that these traits represented a response primarily to levels of larval crowding. Thus, the overall picture that emerged from these two studies was that long-term exposure to larval crowding in *Drosophila* selected for increased larval feeding and foraging activity, at the cost of efficiency at converting food to biomass, a greater tendency to pupate away from the food, and a higher tolerance to toxic levels of metabolic waste. It makes sense that these traits would contribute to greater pre-adult survivorship in a crowded larval culture characterized by diminishing food levels, greater chance of pupal drowning on the increasingly mushy food surface, and a rapid buildup of levels of nitrogenous waste. This has been the canonical view of adaptations to crowding in *Drosophila* for the past fifteen years or so (Mueller 1997; Joshi et al. 2001; Prasad and Joshi 2003; Mueller 2009; Mueller and Cabral 2012).

The notion that faster feeding is a strong correlate of pre-adult competitive ability in *Drosophila* has a lot of support. Populations selected for faster feeding rate were also found to be better competitors (Burnet et al. 1977)), and populations selected for rapid pre-adult development evolved both reduced feeding rate and reduced competitive ability (Prasad et al. 2001; Shakarad et al. 2005; Rajamani et al. 2006). Similarly, populations selected for increased parasitoid resistance evolved both reduced feeding rate and reduced competitive ability (Fellowes et al. 1998, 1999). The notion that there is a cost to faster feeding (Joshi and Mueller 1996) was supported by the rapid return of feeding rates to control levels when CU populations were maintained at moderate densities (Joshi et al. 2003). There is also evidence for a trade-off between urea/ammonia tolerance and larval feeding rate. Populations selected for greater urea and ammonia tolerance, respectively, also showed reduced larval feeding rate (Borash et al. 2000) and larval foraging path length (Mueller et al. 2005), and populations selected for greater urea tolerance did not show higher survivorship than controls at high larval density (Shiotsugu et al. 1997). In general, larval feeding rate and forgaing path length appear to be positively correlated (Joshi and Mueller 1988, 1996; Sokolowski et al. 1997; Borash et al. 2000; Prasad et al. 2001; Mueller et al. 2005). Thus, the evolution of competitive ability in *Drosophila* seems to be the outcome of a balance between mutually antagonistic traits like increased larval feeding and forgaing behaviour (and perhaps pupation height), greater tolerance to nitrogenous wastes, and a reduced efficiency of conversion of food to biomass. Indeed, the CU populations exhibited a temporal polymorphism for two of these traits: offspring of early eclosing flies in a crowded culture showed higher feeding rates, whereas offspring of late eclosing flies showed greater urea/ammonia tolerance than controls (Borash et al. 1998).

Given that the above view of adaptation to larval crowding in *Drosophila* was built around studies on a single species (*D. melanogaster*), we wanted to investigate whether other species of *Drosophila* would also respond to larval crowding by evolving essentially the same set of traits. If the genetic architecture of traits relevant to fitness under larval crowding is reasonably conserved across congeners, then we should see a similar pattern of correlated responses to selection for adaptation to larval crowding in other *Drosophila* species. We report results from two selection experiments, on *D. ananassae* and *D. nasuta nasuta,* involving selection for adaptation to larval crowding. *D. ananassae* is ecologically and phylogenetically closer to *D. melanogaster* (*Sophophora* Subgenus, Melanogaster Group, Melanogaster Subgroup), being a cosmopolitan human commensal and belonging to the Melanogaster Group, Ananassae Subgroup and Ananassae Species Complex, whereas *D. n. nasuta* belongs to the *Drosophila* Subgenus, Immigrans Group, Nasuta Subgroup and Frontal Sheen Complex, and is found primarily in orchards and open land. Our results showed that selected populations of both *D. ananassae* and *D. n. nasuta* exhibited similar patterns of correlated responses to selection for adaptation to larval crowding, but that these populations evolved greater competitive ability through traits very different from those seen earlier in *D. melanogaster* populations subjected to similar selection.

## Materials and methods

### Experimental populations

This study used eight laboratory populations each of *D. ananassae* and *D. n. nasuta.* Four control *D. ananassae* populations (AB_1–4_: **<u>A</u>**nanassae **<u>B</u>**aseline) were derived from a single population, initiated in May-June 2001 with about 300 wild-caught females from Bangalore, India, and maintained as a single population on a 21-day discrete generation cycle for 34 generations (Sharmila Bharathi *et al*. 2003). The four AB populations were maintained on a 21-day discrete generation cycle at 25° ± 1 C, ~90% relative humidity, constant light and on cornmeal medium. Larval density was regulated at 60–80 larvae per vial (9 cm × 2.4 cm) with 6 mL food. Forty vials were set up per replicate population. Twelve days after egg-collection, eclosed adults (1500–1800) from all 40 vials were collected into Plexiglas cages (25 × 20 × 15 cm^3^) containing a Petridish (9 cm diameter) of food that was changed every alternate day, and a moistened ball of cotton. On day 18 after egg-collection, the flies were given food along with a generous smear of live yeast-acetic acid paste. On day 21 from the previous egg-collection, fresh food plates were put into the cages and eggs collected from them after 18 h to initiate the next generation.

From the AB populations, four populations (ACU_1-4_: **<u>A</u>**nanassae **<u>C</u>**rowded as larvae and **<u>U</u>**ncrowded as adults) were derived, one each from each of AB_1-4_, two generations after the AB populations were established. The ACU populations were selected for adaptation to larval crowding by subjecting them to a density of 550–600 eggs per vial with 1.5 mL of food. In initial generations the density was lower; the final densities were attained by generation 15 of ACU selection. The ACU populations were otherwise maintained the same as the AB controls, except that only 20 vials of eggs were collected each generation (to keep the number of breeding adults similar to controls: 1500–1800), and the collection of eclosed adults into the cages continued until day 18 from previous egg-collection, as eclosion in a crowded culture is staggered over several days.

Four control *D. n. nasuta* populations (NB_1-4_: **<u>N</u>**asuta **<u>B</u>**aseline) were derived, after 24 generations as a single population on a 21 day discrete generation cycle, from a laboratory population established using about 70 females collected from orchards and domestic garbage dumps in different parts of Bangalore, India, during October-November 2001 (Sharmila Bharathi *et al.* 2003). The four NB populations were maintained on a 21-day discrete generation cycle at 25° ± 1 C, ~90% relative humidity, constant light and on cornmeal medium. The larval density was regulated at 60–80 larvae per vial (9 cm × 2.4 cm) with 6 mL food. Fifty two vials were set up per replicate population to keep the number of breeding adults at about 1500–1800. Twelve days after egg-collection, eclosed adults from all 40 vials per replicate population were collected into Plexiglas cages (25 × 20 × 15 cm^3^) containing a Petridish of food that was changed every alternate day, and a moistened ball of cotton. On day 18 after egg-collection, the flies were given food along with a generous smear of live yeast-acetic acid paste. On day 21 from the previous egg-collection, fresh food plates were put into the cages and eggs collected from them after 18 h to initiate the next generation.

From the NB populations, four populations (NCU_1-4_: **<u>N</u>**asuta **<u>C</u>**rowded as larvae and **<u>U</u>**ncrowded as adults) were derived, one each from each of NB_1-4_, two generations after the NB populations were established. The NCU populations were selected for adaptation to larval crowding by subjecting them to a density of 350–400 eggs per vial with 2 mL of food. In initial generations the density was lower; the final densities were attained by generation 15 of NCU selection. This difference from ACU in larval density is to compensate for the larger size at each stage of the *D. nasuta* larvae. The NCU populations were otherwise maintained in the same way as the NB controls, with similar adult population size, except that the collection of eclosed adults into the cages continued until day 18 from previous egg-collection, and only 40 vials with eggs were set up.

Since each ACU or NCU population was derived from one control population, selected and control populations of each species bearing identical numerical subscripts are more related to each other than to other populations with which they share the selection regime. Therefore, control and selected populations with identical subscripts were treated as blocks, representing ancestry, in the statistical analyses.

### Collection of flies for assays

All control and selected populations were maintained under common (control-type) rearing conditions for one complete generation prior to assays, to eliminate non-genetic parental effects. The progeny of these flies, hereafter ‘standardized flies’, were then used for the various assays. To obtain progeny for assays, standardized flies in cages were provided yeast-acetic acid paste on food for three days before egg collection. A fresh Petridish with food was then placed in the cages and the flies were allowed to lay eggs for ~14 h, after which eggs were removed from the food with a moistened paintbrush and placed into vials for setting up the various assays. All assays were conducted at 25 ± 1°C, under constant light.

### Pre-adult survivorship

After 42 generations of ACU selection, eggs laid by standardized flies were placed into vials at a density of either 70 or 600 per vial containing 1.5 mL of food. Eight such vials were set up for each replicate AB and ACU population at each density in single-species culture. Eight such vials at each density were also set up in two-species cultures, in competition with a common competitor, a white eyed mutant population of *D. melanogaster*, maintained for about 90 generations in the laboratory on a three week discrete generation cycle. The white eyed population was derived from spontaneously occurring mutant individuals in the JB populations (Sheeba et al. 1998) in our laboratory. For the two-species cultures, eggs laid by standardized ACU, AB or white eyed flies were collected and placed into vials at a density of either 70 (35 ACU or AB eggs and 35 eggs from the white eyed mutant population) or 600 (300 ACU or AB eggs and 300 eggs from the white eyed mutant population) per vial containing 1.5 mL of food. Eight such vials were set up for each replicate AB and ACU population at each density. The number of flies eclosing in each vial was recorded and used to calculate pre-adult survivorship.

At generation 76 of NCU selection, an assay similar to that described above was set up using the NB and NCU populations and the white eyed *D. melanogaster.* The only difference from the *D. ananassae* assay was that the low and high density treatments comprised of 70 or 350 eggs in vials with 2 mL of food.

### Duration of pre-adult life-stages and pre-adult development time

After 53 generations of ACU selection, pre-adult development time was assayed at a high density of 600 eggs per vial with 1.5 mL of food. Eggs from standardized AB and ACU flies were dispensed into each vial using a moistened paintbrush. Eight such vials were set up per replicate population. After the pupae darkened, vials were checked every 6 h and the number of eclosing flies recorded.

After 71 generations of ACU selection, pre-adult development time was assayed at a low density of 30 eggs per vial with 6 mL of food (An earlier study at generation 41 of ACU selection had shown that pre-adult development time did not differ significantly between low densities of 30 or 70 eggs per vial with 6 mL of food: data not shown). Eggs from standardized AB and ACU flies were dispensed into each vial using a moistened paintbrush. Ten vials were set up per replicate population. After the pupae darkened, the vials were monitored for eclosion at 2 h intervals, and the number of eclosing flies recorded. As part of the same assay, egg hatching time and the duration of each larval instar and the pupal stage were also determined. For assaying egg hatching time, 30 eggs from the standardized flies were arranged on a small agar cube in a food vial in six rows of five eggs each. Ten such vials were set up per population. Fifteen hours after egg laying, the vials were checked for any hatched eggs once every hour, till no eggs hatched for three consecutive hours. For assaying instar and pupal duration, eggs of approximately identical age were harvested over a three hour period from the standardized flies and dispensed into vials with 6 mL of food at a density of 30 eggs per vial. Sixty such vials were set up per population. Forty three hours after the midpoint of the three hour egg laying period, four vials per population were removed from the incubators and immersed in hot water. The dead larvae were removed and kept in 70% ethanol for subsequent examination. Every two hours, this process was repeated. The larval instars were differentiated based on the number of ‘teeth’ in the larval mouth hooks. From these data, the number of larvae of each instar present in each two-hourly sample was determined, and the median time of each molt was obtained by interpolation. For pupal duration and pre-adult development time, ten vials per population were set up with 30 eggs in 6 mL of food. After the first pupa was seen, the vials were screened every two hours and any new pupae that had formed were marked on the vial with different coloured marker pens. Thereafter, the vials were monitored for eclosion and the number of eclosing males and females in each vial was determined every two hours. These observations yielded data on egg to pupa and egg to adult development time, from which the pupal duration could be calculated.

After 84 generations of NCU selection, pre-adult development time was assayed at a high density of 350 eggs per vial with 2 mL of food. Eggs from standardized NB and NCU flies were dispensed into each vial using a moistened paintbrush. Eight such vials were set up per population. After the pupae darkened, the vials were checked every 6 h and the number of eclosing flies recorded.

After 62 generations of NCU selection, egg hatching time, larval instar duration, pupal duration and pre-adult development time of the NB and NCU populations were assayed exactly as described above for the ACU and AB populations.

### Larval feeding rate

After 71 generations of ACU selection, the feeding rates of AB and ACU larvae were measured at physiologically equalized ages, based on the difference in AB and ACU development time. This was done by collecting eggs from the standardized ACU flies 5 h later than the AB flies. Thus, at the time of assay, ACU larvae were 58 h old, whereas AB larvae were 63 h old and, thus, approximately in the same relative stage of their larval development. Following Joshi and Mueller (1996), about a hundred eggs laid over a four hour period were collected from standardized flies and placed into two Petridishes with non-nutritive agar each for AB and ACU populations. Twenty-four hours later, twenty-five newly hatched larvae were transferred from these agar Petridishes to a Petridish containing a thin layer of non-nutritive agar overlaid with 1.5 mL of 37.5% yeast suspension. Four such Petridishes were set up per population. The larvae were then allowed to feed for 58 (ACU) or 63 (AB) h, by which time they were in the early third instar. At this point, 20 larvae from each population were assayed for feeding rate, following the procedure of Joshi and Mueller (1996), by placing them individually in a small Petridish (5 cm diameter) containing a thin layer of agar overlaid with a thin layer of 10% yeast suspension. After allowing for a 15 sec acclimation period, feeding rate was measured under a stereozoom microscope as the number of cephalopharyngeal sclerite retractions in a 1 min period. Selected and control populations, matched by the subscripted indices, were assayed together, with one larva from the selected population and one from the control population being assayed alternately. The same procedure was followed for assaying larval feeding rate of 20 larvae from each NCU and NB population, after 77 generations of NCU selection.

### Larval foraging path length

Twenty early third instar larvae for each population were assayed for larval foraging path length. The collection of larvae for the assay was exactly as described above for larval feeding rate assays. For assaying foraging path length, individual larvae were placed in a small Petridish (5 cm diameter) containing a thin layer of agar overlaid with a thin layer of 10% yeast suspension. After allowing for a 15 sec acclimation period, the larvae were allowed to move around on the Petridish for 1 min. The path traversed by the larvae on the yeast surface was traced onto a transparency sheet and later measured with a thread and ruler. Selected and control populations, matched by subscripted indices, were assayed together at generations 52 and 81 of ACU and NCU selection, respectively.

### Pupation height

Eggs from standardized flies were collected and placed into vials with 6 mL of food at a density of 50 (ACU/AB) or 70 (NCU/NB) eggs per vial. Ten and eight such vials were set up per population for *D. ananassae* and *D. n. nasuta,* respectively. Once all pupae had formed, pupation height of each pupa was measured, following Joshi and Mueller (1993), as the distance between the surface of the food medium to the point between the anterior spiracles of the pupa. Any pupa on or touching the surface of the food was given a pupation height of zero. Assays were conducted at generations 39 and 34 of ACU and NCU selection, respectively.

### Larval urea and ammonia tolerance

After 49 generations of ACU selection, eggs laid over a four hour period by standardized flies were collected and exactly 30 eggs per 6 mL food were placed into vials at three concentrations each of urea (0, 14 and 18 g/L) or ammonia (0, 15 and 30 g/L NH_4_Cl) in the food. These are values that allowed the detection of differences in urea/ammonia tolerance between selected and control populations in previous studies on *D. melanogaster* (Shiotsugu et al. 1997; Borash et al. 1998). Ten vials were set up per population at each concentration of either urea or ammonia, and the number of eclosing flies in each vial was recorded to calculate pre-adult survivorship. Both urea and ammonia were used as there is some confusion about which compound actually increases in concentration in crowded larval cultures, although it is likely that it is ammonia in the case of *D. melanogaster* (Botella et al. 1985; Borash et al. 1998). The NCU and NB populations were assayed in an identical manner after 76 generations of NCU selection, except that the egg-laying period was 12 hours, and the concentrations used were 0, 9 and 11 g/L for urea and 0, 15 and 20 g/L for ammonia, as earlier studies had shown that *D. n. nasuta* larvae were more sensitive to urea and ammonia than their *D. ananassae* counterparts, and *D. n. nasuta* females are not as fecund as *D. ananassae* females (data not shown).

### Minimum feeding time and dry weight after minimum feeding

These assays were conducted only on the ACU and AB populations. After 45 generations of ACU selection, critical minimum feeding time for pupation was assayed by setting up freshly hatched larvae, from eggs laid by standardized flies, onto Petridishes with agar overlaid with 1.5 mL of 37.5% yeast suspension, as described above for the feeding rate and foraging path length assays. Twenty five larvae were placed into each Petridish, and sixty such Petridishes were set up per population. At 46, 49, 52 and 55 h after egg hatch, a total of 150 larvae per population per time point were removed from the yeast, gently washed in water to remove any yeast sticking to their bodies, and placed in ten vials containing 5 mL of non-nutritive agar, at a density of 15 larvae per vial. These vials were subsequently monitored for pupation and eclosion, and the pre-adult survivorship after feeding for different periods of time noted. Using the information from this assay, a second assay was carried out after 68 generations of ACU selection to measure the dry weights at eclosion of ACU and AB flies that had fed as larvae for different durations of time, roughly corresponding to pre-adult survivorship of 13, 25, 50 and 60%, respectively (feeding for 54, 56, 59 and 62 h for AB, and 50, 55, 58 and 60 h for ACU populations, respectively). Freshly hatched larvae were collected, shifted to yeast and then removed and placed into agar vials after feeding for different durations corresponding to pre-adult survivorship of 13, 25, 50 and 60% for ACU and AB populations, exactly as described above. Once eclosions began in the agar vials, flies were collected every 4 h and frozen for subsequent weighing. Frozen flies were sorted into batches of five males or five females each and dried at 70°C for 36 h before weighing in batches.

### Statistical analyses

Since assays on ACU/AB and NCU/NB populations were conducted at different times, data from the two species were analyzed separately. All traits were subjected to completely randomized block analyses of variance (ANOVA). All ANOVAs treated replicate population 1..4 (representing ancestry) as random blocks crossed with the fixed factor selection regime. Additional fixed factors, crossed with both selection regime and block, were included when relevant. These factors were larval density and type of culture (single- or two-species) for pre-adult survivorship, pre-adult life-stage for life-stage duration, urea or ammonia concentration in the food for urea and ammonia tolerance, and larval feeding duration or survivorship for the assays on minimum critical feeding time and of dry weight at eclosion after feeding for different time durations corresponding to four different pre-adult survivorship levels. Development time data from low and high density were analyzed separately, rather than incorporating larval density as a factor, because the low and high density assays were conducted at different times for each species. Data on pre-adult survivorship were arcsine-squareroot transformed before ANOVA. For all traits, ANOVAs were done on replicate population mean values and, therefore, only fixed factor effects and interactions could be tested for significance. All analyses were implemented using Statistica for Windows rel.5.0 B, (Stat Soft 1995). Multiple comparisons were done using Tukey’s HSD test.

## Results

### Pre-adult survivorship

Pre-adult survivorship at high larval density is the primary trait expected to be under direct selection in populations exposed to high larval crowding each generation. With regard to survivorship at low versus high larval density, the broad pattern of results was very similar in the two species (Figure 1). In both species, pre-adult survivorship was, on an average, significantly higher at low rather than high larval density, and in selected compared to control populations (Figure 1, Table 1). The trend was for survivorship to be higher, on average, in single-species cultures than in two-species cultures, but the difference was significant only in the case of *D. ananassae* (Table 1). Both species showed a significant selection regime × larval density interaction (Table 1), with the survivorship of selected and control populations not differing significantly at low density, and with selected populations showing significantly higher survivorship than controls at high density (Figure 1). Both species also showed a significant culture type × larval density interaction (Table 1). In *D. ananassae,* averaged over selection regimes, survivorship at low density in both types of cultures (single- or two-species) was similar, whereas survivorship at high density was significantly greater in single-species than two-species cultures (Figure 1). In *D. n. nasuta,* the opposite pattern was seen. Averaged over selection regime, survivorship at low density in single-species cultures was higher than in two-species cultures (Figure 1), but not significantly so. However, at high density, survivorship in two-species cultures was higher than in single-species cultures but, again, not significantly so. In fact, the only clear significant difference in the multiple comparisons was that between single-species cultures at low and high densities (Figure 1.). In the case of *D. ananassae,* there was also a significant culture type × selection regime interaction (Table 1), but the only clearly significant differences in multiple comparisons were between survivorship in two-species cultures at high density on the one hand, and in the other three culture type × selection regime combinations on the other (Figure 1).

**Figure 1.**
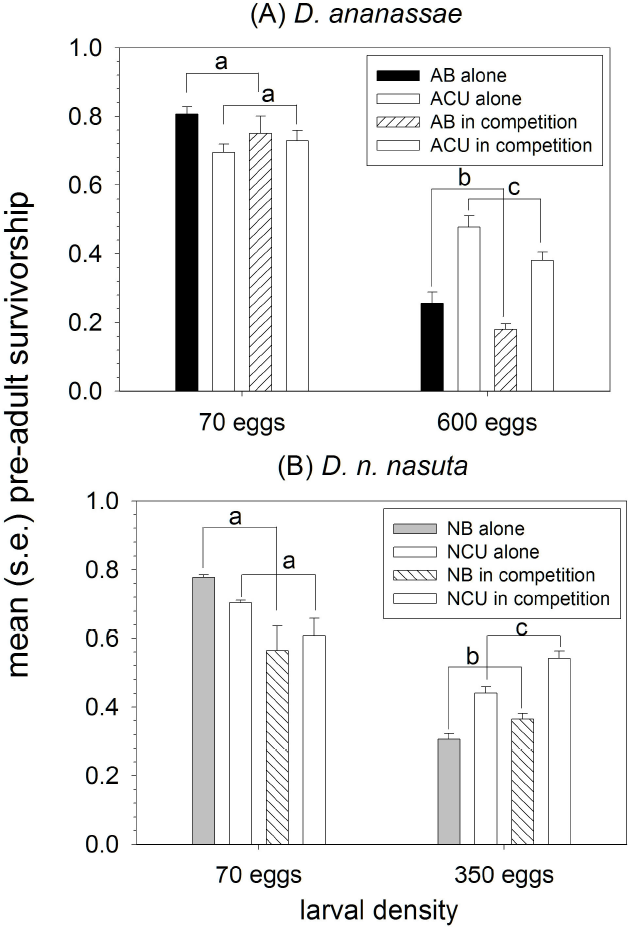
Mean pre-adult survivorship at low and high larval density in the selected and control populations of the two species, when cultured alone or in the presence of equal numbers of eggs of white-eyed *D. melanogaster* competitors. Error bars are standard errors around the mean of the four replicate population mean values. Significant differences between pairs of means are indicated by labeling them with different letters. For both species, as the three-way interaction between type of culture, selection regime and larval density was not significant, only the difference between the mean pre-adult survivorships of selected and control populations at high and low larval densities, averaged over type of culture (single- or two-species), is highlighted.

**Table 1.** Summary of results of four-way ANOVA on mean arcsin squareroot transformed pre-adult survivorship at low and high density in the two sets of selected populations and controls, in single-species cultures and in two-species competitive cultures with equal numbers of white-eye *D. melanogaster* eggs. Since the analysis was done on population means, block and interactions involving block were not tested for significance.

| Effect | ACU/AB |  | NCU/NB |  |
| --- | --- | --- | --- | --- |
| | $F_{1,3}$ | $P$ | $F_{1,3}$ | $P$ |
| culture type | 17.44 | 0.03 | 1.54 | 0.30 |
| selection | 14.11 | 0.033 | 43.67 | <0.01 |
| density | 1621.07 | <0.01 | 51.54 | <0.01 |
| culture type × selection | 11.81 | 0.04 | 5.50 | 0.10 |
| culture type × density | 10.03 | 0.05 | 17.46 | 0.03 |
| selection × density | 525.03 | <0.01 | 60.88 | <0.01 |
| culture type × selection × density | 2.94 | 0.19 | 2.12 | 0.24 |

### Pre-adult development time

In general, pre-adult development time in *D. n. nasuta* was higher than that in *D. ananassae,* as also noted earlier (Sharmila Bharathi et al. 2004). Pre-adult development time was, in general, greater in males than in females, and greater at high rather than low larval density, as is usually the case in *Drosophila* (Figures 2, 3, Table 2).

**Figure 2.**
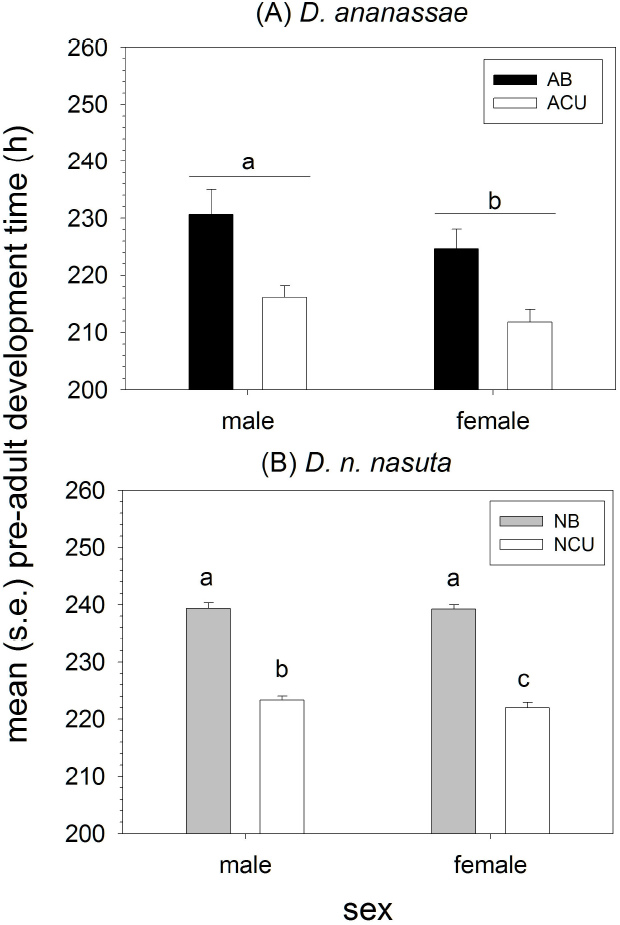
Mean male and female pre-adult development time at low larval density in the selected and control populations of the two species. Error bars are standard errors around the mean of the four replicate population mean values. For both species, significant differences between pairs of means are indicated by labeling them with different letters. For *D. ananassae*, as the interaction between sex and selection regime was not significant, only the difference between the mean pre-adult development time of males and females, averaged over selection regimes is highlighted.

**Figure 3.**
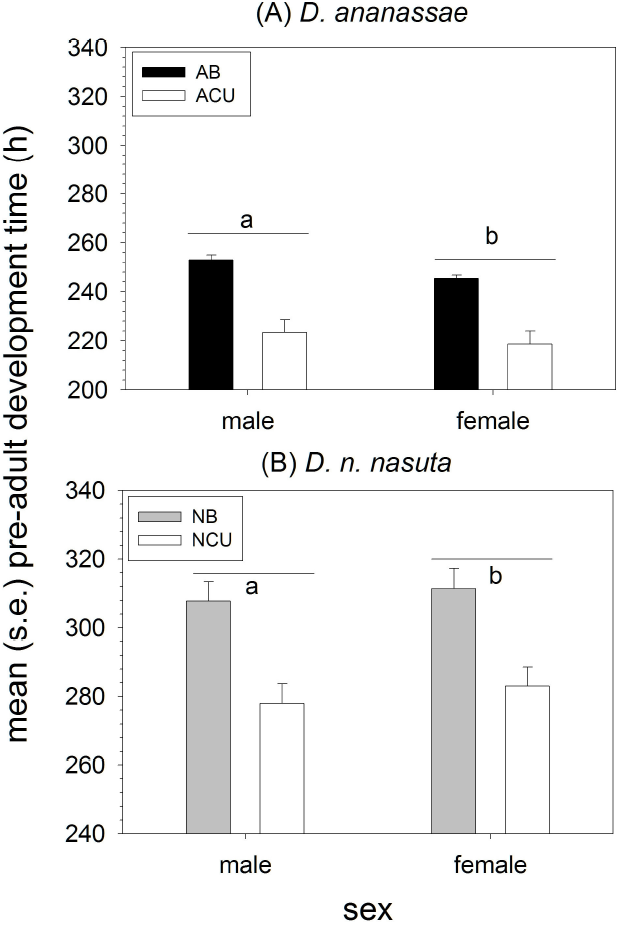
Mean male and female pre-adult development time at high larval density in the selected and control populations of the two species. Error bars are standard errors around the mean of the four replicate population mean values. Significant differences between pairs of means are indicated by labeling them with different letters. For both species, as the interaction between sex and selection regime was not significant, only the difference between the mean pre-adult development time of males and females, averaged over selection regimes is highlighted.

**Table 2.** Summary of results of two-way ANOVA on mean pre-adult development time, assayed at low and high larval density at different generations in the two sets of selected populations and controls. Since the analysis was done on population means, block and interactions involving block were not tested for significance.

|  | <u>ACU/AB</u> |  | <u>NCU/NB</u> |  |
| --- | --- | --- | --- | --- |
| <u>Low density</u> |  |  |  |  |
| Effect | $F_{1,3}$ | $P$ | $F_{1,3}$ | $P$ |
| selection | 49.23 | <0.01 | 115.50 | <0.01 |
| sex | 82.32 | <0.01 | 3.75 | 0.15 |
| selection × sex | 1.93 | 0.26 | 18.61 | 0.02 |
| <u>High density</u> |  |  |  |  |
| Effect | $F_{1,3}$ | $P$ | $F_{1,3}$ | $P$ |
| selection | 60.37 | <0.01 | 364.65 | <0.01 |
| sex | 41.18 | <0.01 | 13.09 | 0.04 |
| selection × sex | 8.58 | 0.06 | 5.49 | 0.10 |

Interestingly, in both species at both larval densities, there was a significant main effect of selection regime, with selected populations consistently showing substantially lower development time than control populations at both densities (Figures 2, 3, Table 2). On an average, AB development time was higher than that of ACU populations by about 14 and 29 h at low and high larval density, respectively (Figures 2A, 3A), whereas that of NB populations was greater than that of NCU populations by about 17 and 29 h at low and high larval density, respectively (Figures 2B, 3B). In three of the four assays, there was no significant selection regime × sex interaction (Table 2). In *D. n. nasuta* at high larval density, control males and females had similar development times whereas NCU males had significantly higher development time than NCU females, albeit by just about 1 h (Figure 3B).

### Pre-adult life-stage duration

The overall pattern of pre-adult life-stage durations in both species was similar to what is seen in *Drosophila* spp. in general. Consistent with the pre-adult development time results, both species showed a main effect of selection regime (Table 3), with life-stage durations being greater in control rather than selected populations, on an average (Figure 4). In *D. ananassae*, a 20 h egg stage was followed by two larval instars of about 24 h each, a third larval instar of about 46 h and a pupal stage lasting about 87 h (Figure 4A); there was no significant selection regime × life-stage duration interaction. In *D. n. nasuta,* all pre-adult life-stages differed significantly from one another in duration, with a 23 h egg stage, followed by two larval instars of about 27 and 29 h, respectively, a longer third larval instar of about 63 h and a pupal stage lasting about 87 h (Figure 4B). The longer larval instar durations in *D. n. nasuta* are consistent with their larger adult size compared to *D. ananassae* (Sharmila Bharathi et al. 2004). *D. n. nasuta* also showed a significant selection regime × life-stage duration interaction (Table 3), with a large (~12 h) reduction in third instar duration in the NCU populations (Figure 4B). We note that, given the time interval between subsequent observations (1 h for egg-hatch; 2 h for larval instars), our screening might be too coarse to pick up small but consistent differences on the order of 30 min – 1 h. However, the differences between selected and control populations in the durations of different pre-adult life-stages, though often not significant, did add up roughly to the overall difference seen in pre-adult development time between selected and control populations in both species.

**Figure 4.**
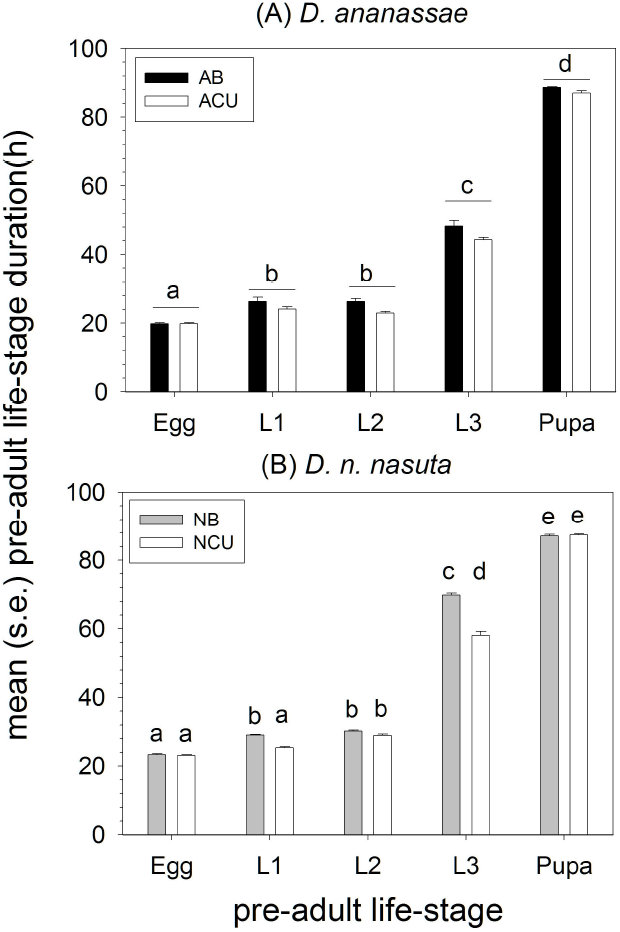
Mean duration in hours of different pre-adult life-stages at low larval density in the selected and control populations of the two species. Error bars are standard errors around the mean of the four replicate population mean values. For each species, significant differences between pairs of means are indicated by labeling them with different letters. For *D. ananassae,* the interaction between selection regime and life-stage was not significant. Consequently, only the differences in mean duration between life-stages, averaged over block and selection regime, have been highlighted.

**Table 3.** Summary of results of three way ANOVA on mean pre-adult lifestage (egg, L1, L2, L3, pupa) duration in the two sets of selected populations and controls. Since the analysis was done on population means, block and interactions involving block were not tested for significance.

|  | <u>ACU/AB</u> |  | <u>NCU/NB</u> |  |
| --- | --- | --- | --- | --- |
| Effect | $F_{1,3 \text{ sel/ } 4,12 \text{ else}}$ | $P$ | $F_{1,3 \text{ sel/ } 4,12 \text{ else}}$ | $P$ |
| selection | 168.38 | <0.01 | 115.50 | <0.01 |
| life-stage | 1778.55 | <0.01 | 9485.70 | <0.01 |
| selection × life-stage | 1.46 | 0.28 | 37.05 | <0.01 |

### Larval traits

Surprisingly, larval feeding rate did not differ significantly between selected and control populations in either species (Table 4). Mean (± s.e.) larval feeding rates in sclerite retractions per min were, in fact, extremely similar between selected and control populations of both species (AB: 128.33 ± 3.13; ACU: 131.55 ± 3.05; NB: 112.76 ± 5.29; NCU: 113.03 ± 4.34). In both species, selected populations showed significantly greater mean (± s.e.) pupation height in cm than control populations (AB: 1.02 ± 0.28; ACU: 1.60 ± 0.20; NB: 0.44 ± 0.12; NCU: 1.54 ± 0.11) (Table 4). In case of larval foraging path length, the results varied between species, with *D. ananassae* showing a significant main effect of selection regime while *D. n. nasuta* did not (Table 4). AB populations had mean (± s.e.) foraging path length in cm of 5.52 ± 0.61, compared to 7.38 ± 0.60 in the ACU populations. NB and NCU populations had mean (± s.e.) foraging path length in cm of 4.24 ± 0.53 and 4.08 ± 0.43, respectively.

**Table 4.**
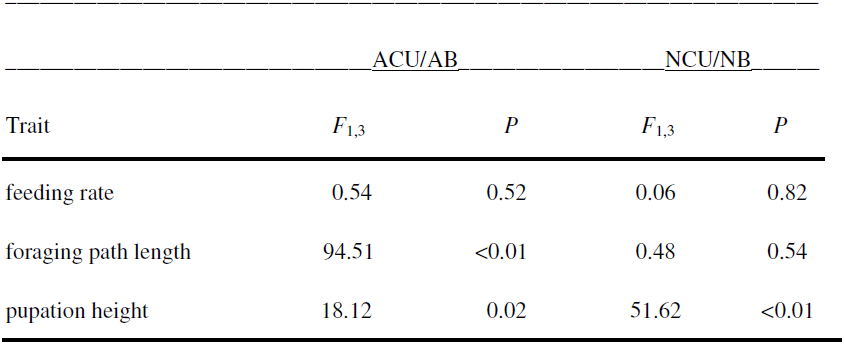
Summary of results of two way ANOVA on mean larval feeding rate, foraging path length and pupation height in the two sets of selected populations and controls. Since the analysis was done on population means, block and interactions involving block were not tested for significance; only the *F* and *P* values for the main effect of selection regime are shown.

### Larval urea and ammonia tolerance

There was no clear pattern to the results of assays on pre-adult survivorship in the presence of metabolic wastes like urea (Figure 5) and ammonia (Figure 6) in the food medium, except for strong evidence for the toxic effects of these compounds, reflected in significant main effects of concentration in all four ANOVAs (Table 5).

**Figure 5.**
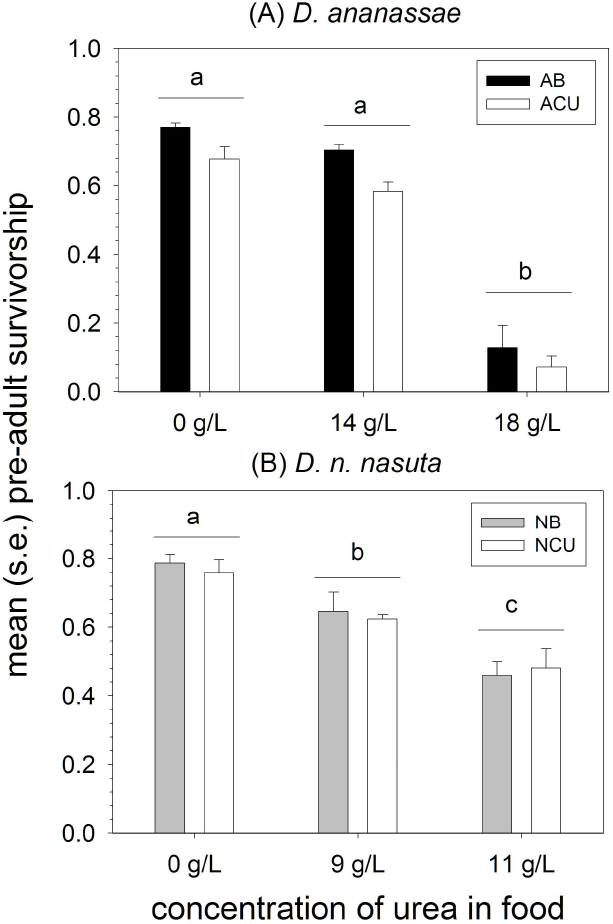
Mean pre-adult survivorship at low density with three different levels of urea in the food medium in the selected and control populations of the two species. Error bars are standard errors around the mean of the four replicate population mean values. For each species, significant differences between pairs of means are indicated by labeling them with different letters. For both species, the interaction between selection regime and urea level was not significant. Consequently, only the differences between mean survivorship at different urea levels, averaged over block and selection regime, have been highlighted.

**Figure 6.**
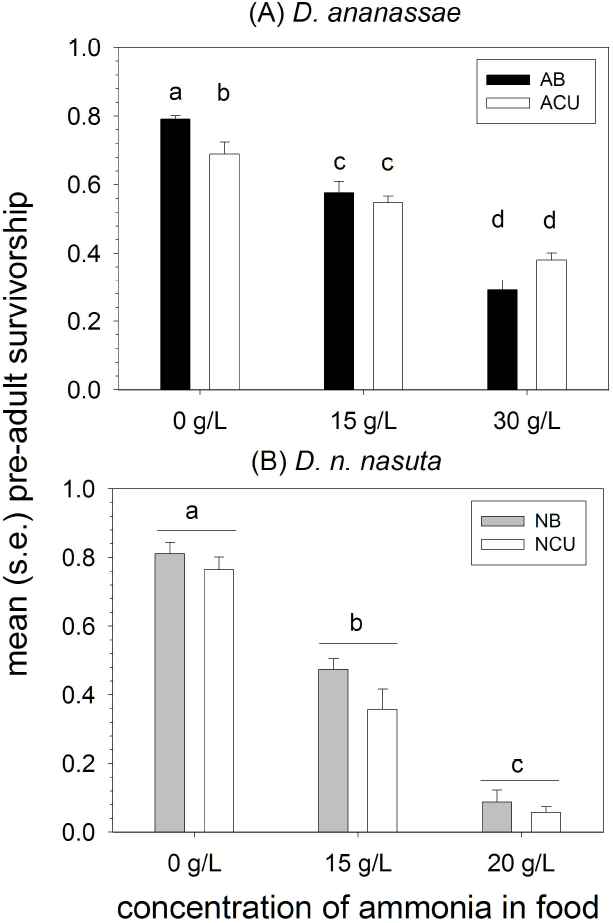
Mean pre-adult survivorship at low density with three different levels of ammonia in the food medium in the selected and control populations of the two species. Error bars are standard errors around the mean of the four replicate population mean values. For each species, significant differences between pairs of means are indicated by labeling them with different letters. For *D. n. nasuta* the interaction between selection regime and ammonia level was not significant. Consequently, only the differences between mean survivorship at different ammonia levels, averaged over block and selection regime, have been highlighted.

**Table 5.**
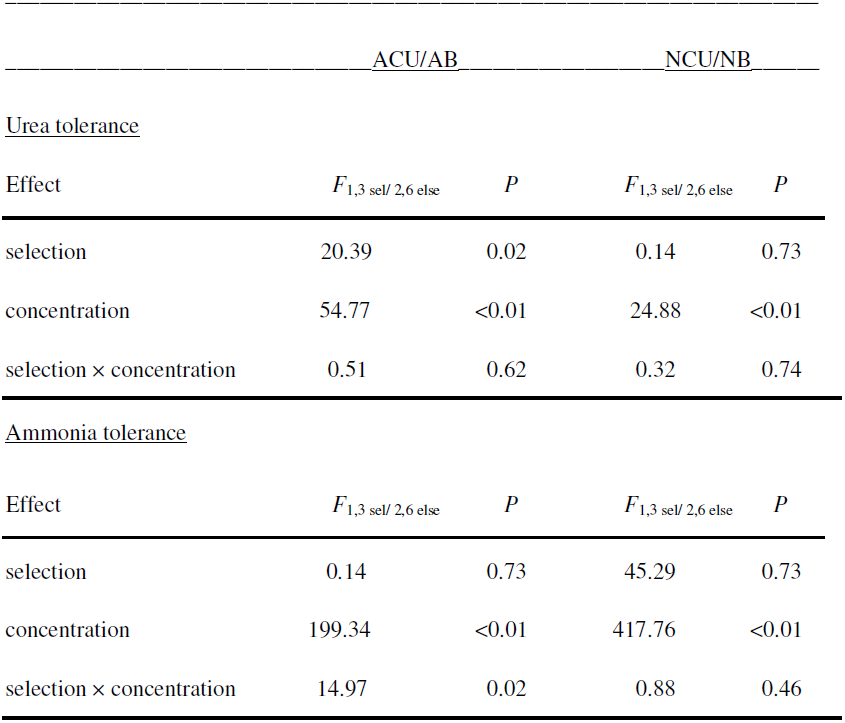
Summary of results of three way ANOVA on mean arcsin squareroot transformed pre-adult survivorship at three different concentrations of urea or ammonia in the food medium in the two sets of selected populations and controls. Since the analysis was done on population means, block and interactions involving block were not tested for significance.

ACU populations had lower survivorship than AB controls across all levels of urea, including 0 g/L, and thus showed no evidence of greater or lesser urea tolerance than the AB controls (Figure 5A), with a significant main effect of selection regime but no significant selection regime × concentration interaction (Table 5). There was suggestive evidence for greater ammonia tolerance in the ACU compared to the AB populations (Figure 6A). At 0 g/L of ammonia in the food, ACU populations had significantly lower survivorship than AB controls, whereas at 30 g/L, the ACU survivorship was higher, though not significantly so, than the AB survivorship (Figure 6A). The selection regime × concentration interaction was also significant for this assay (Table 5).

In the case of *D. n. nasuta,* there was no significant effect of selection regime or of the selection regime × concentration interaction in the urea tolerance assay (Table 5). However, there was a non-significant trend of NCU survivorship being lower than NB at 0 g/L and higher than NB at 11 g/L (Figure 5B). In case of ammonia, NCU populations had consistently lower survivorship than NB controls at all concentrations (Figure 6B) and there was no significant effect of selection regime or of the selection regime × concentration interaction, yielding no suggestion of increased ammonia tolerance in the NCU populations.

### Minimum critical feeding time

In the minimum critical feeding time assay, only significant main effects of selection regime and feeding duration were seen (Table 6), with pre-adult survivorship increasing from feeding durations from 46–55 h, and with ACU survivorship being higher than that of the AB populations at every feeding duration (Figure 7A). At 55 h of feeding as larvae, ACU survivorship was over 50% while that of the AB populations was less than 30% (Figure 7A). Overall, ACU populations seemed to attain similar levels of survivorship approximately 6 h before the AB controls (Figure 7A), which is commensurate with the 54.8 h difference between their first and second instar durations combined. These results suggest that larvae in the ACU populations attain their minimum critical size for pupation approximately 5–6 h before their AB controls.

**Table 6.**
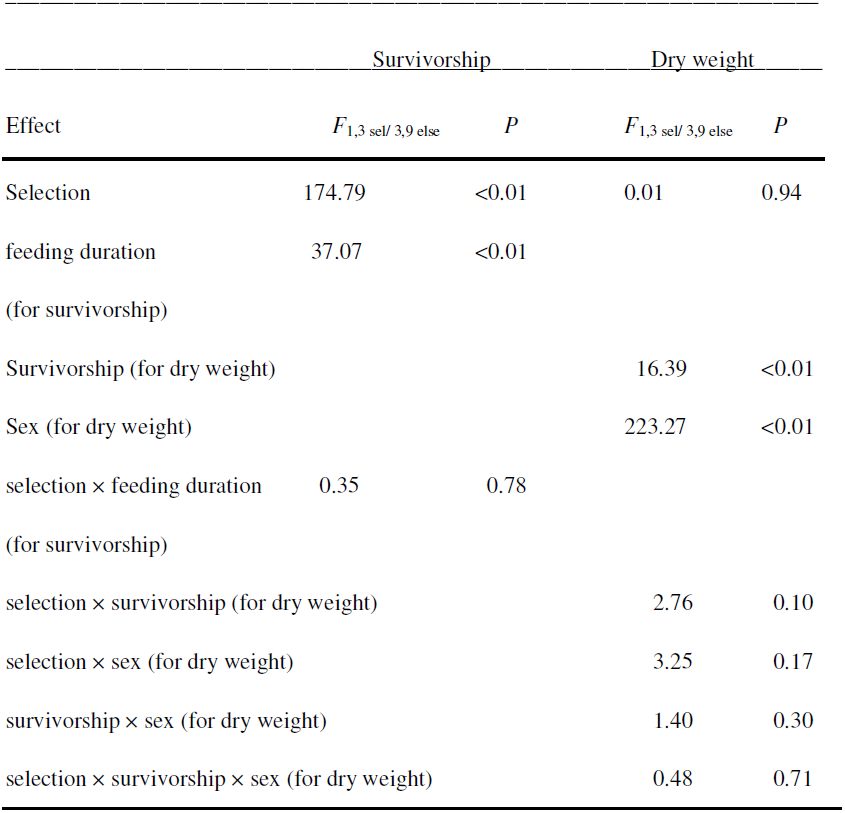
Summary of results of three way ANOVA on mean arcisn squareroot transformed pre-adult survivorship (factors: block, selection regime and larval feeding duration) and mean dry weight (factors: block, selection regime, sex and pre-adult survivorship) of eclosing flies after larvae were allowed to feed for different time periods in the AB and ACU populations. Since the analysis was done on population means, block and interactions involving block were not tested for significance.

**Figure 7.**
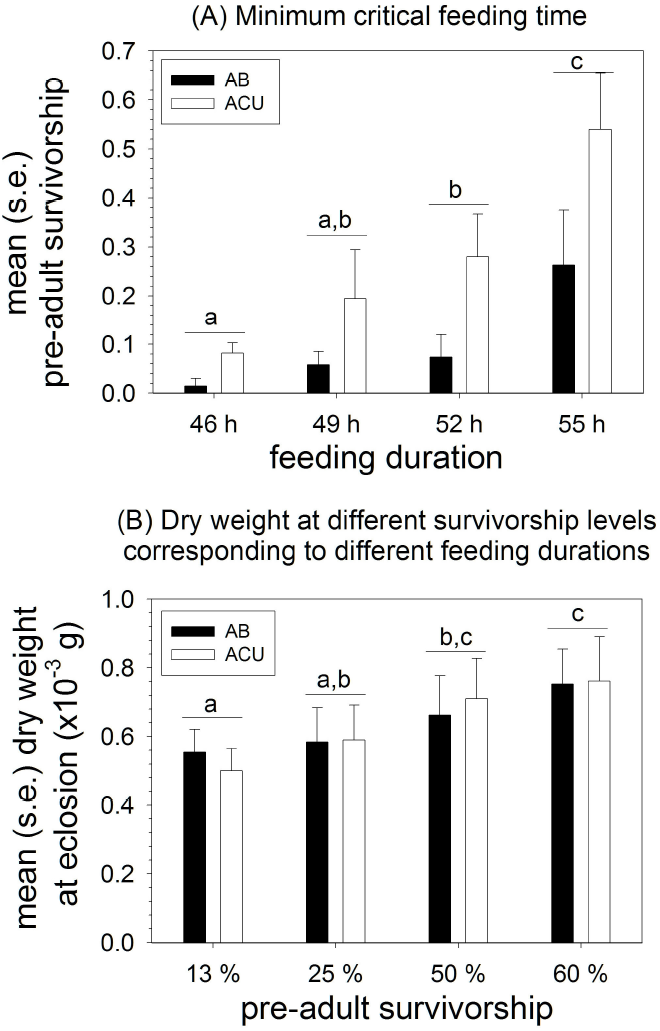
(A) Mean pre-adult survivorship after larval feeding for different amounts of time, and (B) mean dry weight (in 10^−3^ g) at eclosion after larval feeding for different amounts of time corresponding to four different mean pre-adult survivorship levels, in the *D. ananassae* selected and control populations. Error bars are standard errors around the mean of the four replicate population mean values. For both traits, significant differences between pairs of means are indicated by labeling them with different letters. For both traits, the interaction between selection regime and feeding duration/survivorship was not significant. Consequently, only the differences in mean survivorship between the different levels of feeding duration, and differences in mean dry weight at eclosion at different levels of survivorship, averaged over block, selection regime and sex, have been highlighted.

In the ANOVA on data from the assay of dry weight at eclosion of individuals that had fed for different durations of time as larvae, corresponding to four different levels of pre-adult survivorship, only the main effects of sex and survivorship were significant (Table 6). On an average, females were significantly heavier than males, and dry weight at eclosion tended to increase with survivorship (Figure 7B). At no survivorship level was there a significant difference between the dry weights of AB and ACU flies (Figure 7B), suggesting that the minimum critical size for pupation in the selected and control populations is probably not different, even though the ACU populations attain it faster than the AB controls.

### Discussion

It is clear that both the ACU and NCU populations did evolve adaptations to larval crowding over the course of selection: in both species, pre-adult survivorship at high density and competitive ability was greater in selected than in control populations (Figure 1, Table 1). The overall pattern of correlated responses to selection in the two species was also similar, and, more importantly, different from that reported earlier in case of *D*. *melanogaster* in that it did not involve the evolution of higher feeding rates (Table 4).

In clear contrast to what was seen earlier in *D. melanoagster,* both ACU and NCU populations evolved faster pre-adult development time than their controls (Table 2), with the difference being apparent at both low (Figure 2) and high (Figure 3) assay density. The pre-adult development time difference between selected and control populations is entirely due to reduction in the duration of larval instars (Figure 4). In absolute terms, the reductions in pre-adult development time in the selected populations (~14 h in ACU, ~17 h in NCU) are quite large. For comparison, populations of *D. melanogaster* subjected to strong directional selection for reduced pre-adult development time showed a reduction of ~16 h in the larval duration, and ~10 h in pupal duration, after 50 generations of selection (Prasad et al. 2001). In the earlier studies on *D. melanogaster,* CU populations showed no difference from controls in development time at low density, whereas they were faster developing than controls when assayed at high density (Borash and Ho 2001). While the *K*-populations did show faster development than *r*-populations when assayed at low density (Bierbaum et al. 1989), this cannot be unequivocally ascribed to density-dependent selection. The *K*-populations were on an overlapping generation maintenance regime and, relative to a discrete generation regime like that of the *r*-populations, this itself would impose direct selection for faster development. If we put the results on life-stage duration (Figure 4) and development time (Figures 2, 3) together with those from the minimum feeding time assay (Figure 7), the ACU and NCU populations have evolved faster development, especially in the first two larval instars, enabling them to reach the critical minimum size for pupation about 6 h earlier than controls. This is particularly impressive considering that *D. melanogaster* populations selected directly for rapid development reached the critical size only about 2 h before controls after 50 generations of selection (Prasad et al. 2001). At the same time, the weights of ACU and NCU flies are not different from controls after feeding for different durations close to the minimum feeding time (Figure 7, Table 6). Thus, in terms of time, ACU and NCU larvae are clearly more efficient at converting food to biomass than controls during the first two (and an early part of the third) larval instars, becoming equally heavy adults as controls after feeding for about 6 h less. Of course, whether they are more efficient also in terms of food consumed cannot be determined from these experiments. However, these results do suggest that unlike the *K-* and CU populations, (Mueller 1990; Joshi and Mueller 1996), the ACU and NCU populations have evolved to become more rather than less efficient than controls. The faster development of the ACU and NCU populations, relative to controls, is also interesting in the context of observations that faster development correlates with greater competitive ability across species of *Drosophila* (Krijger et al. 2001), even though direct selection for rapid development leads to the evolution of reduced competitive ability (Shakarad et al. 2005) through reduced larval feeding rate, foraging path length and pupation height (Prasad et al. 2001) as well as reduced larval urea tolerance (Joshi et al. 2001).

Unlike in *D. melanogaster*, larval feeding rates did not evolve to become greater in the ACU and NCU populations (Table 4), although these populations clearly evolved greater competitive ability than controls (Figure 1). In fact, we measured feeding rates at various points during ACU and NCU selection, and also at various larval stages, and there was never any difference between selected and control populations in either species (data not shown). Interestingly, larval foraging path length was greater in ACU populations than in AB populations, whereas NCU and NB populations had very similar foraging path lengths (Table 4). Thus, at least in *D. ananassae,* the consistent positive relationship between larval feeding rate and foraging path length seen in *D. melanogaster* (Joshi and Mueller 1988, 1996; Sokolowski et al. 1997; Borash et al. 2000; Prasad et al. 2001; Mueller et al. 2005) seems to be uncoupled. Pupation height evolved to become greater in both ACU and NCU populations, relative to controls, mirroring results from the *K*-populations (Mueller and Sweet 1986), but not the CU populations (Joshi and Mueller 1996; but see also Mueller et al. 1993 and Joshi et al. 2003). One possible reason for the evolution of increased pupation height in the *K*-populations but not the CU populations has been speculated to be the greater hardness and dryness of the cornmeal based food (*r*-and *K*-populations) compared to banana based food (UU and CU populations). It was suggested that banana food being much softer and more fluid compared to cornmeal food, perhaps the UU populations were also under selection for increased pupation height due to greater risk of pupal drowning on the food surface even at their relatively low rearing density (Joshi et al. 2003). In this context, we note that our ACU/AB and NCU/NB populations are maintained on a relatively hard and dry cornmeal food and this might be the reason for the evolution of greater pupation height in selected populations, relative to controls.

Compared to other traits, the results on urea and ammonia tolerance in the selected and control populations were not very clear. While pre-adult survivorship clearly decreased with increasing amounts of urea or ammonia in the food (Figures 5, 6, Table 5), there was no clear evidence for greater urea or ammonia tolerance in the selected populations, except for greater ammonia tolerance in the ACU populations, relative to the AB controls (Figure 6A). There was a slight suggestion of a trend towards increased urea tolerance in the NCU populations, relative to NB controls in that NCU survivorship at 0 g/L was below that of NB populations and became slightly higher than NB populations at 11 g/L (Figure 5B). However, there was no significant selection regime × urea concentration interaction (Table 5), indicating that this is at best a suggestive result. It might be that crowded *D. ananassae* and *D. n. nasuta* cultures have experienced differential levels of urea versus ammonia build-up to which they have adapted. Moreover, we have seen, in other studies with *D. melanogaster*, that the detection of between-population differences in urea/ammonia tolerance is often affected by an interaction between the specific concentrations used and the larval density (Avani Mital, Gitanjali P. Vaidya and Amitabh Joshi, *unpubl. data*). It is, therefore, possible that we were unable to detect significant differences between selected and control populations due to the specific concentrations and larval densities used in our assays of urea and ammonia tolerance. It is also known that larval exposure to urea markedly reduces subsequent fecundity in *D. melanogaster*, and that populations selected for increased larval urea tolerance undergo smaller fecundity declines than controls when reared as larvae on food with urea (Shiotsugu et al. 1997). Thus, it is also possible that the ACU and NCU populations might have evolved greater tolerance to the detrimental effects of urea on fecundity; our experiments did not explore this possibility.

Overall, the crowded *D. ananassae* and *D. n. nasuta* populations seem to have evolved greater competitive ability and pre-adult survivorship at high density primarily through a combination of reduced duration of the larval stage, faster attainment of minimum critical size for pupation, greater efficiency of food conversion to biomass, increased pupation height and, perhaps, greater urea/ammonia tolerance. This is in contrast to *D. melanogaster*, in which crowding adapted populations evolve greater competitive ability and pre-adult survivorship at high density primarily through a combination of increased larval feeding rate and foraging path length, at the cost of reduced efficiency of food conversion to biomass, and greater urea/ammonia tolerance (Mueller 1997; Prasad and Joshi 2003; Mueller 2009). Thus, the ACU and NCU populations appear to have responded to crowding in a manner closer to the canonical notion of *K*-selection, in contrast to the primarily *α*-selection responses shown by the *K*-and CU populations of *D. melanogaster* (see also Dey et al. 2012). Thus, the results underscore the fact that there are, in principle, multiple routes to the evolution of greater competitive ability (Joshi et al. 2001; Dey et al. 2012), something that has also been experimentally demonstrated in the context of inter-specific competition in *Drosophila* (Joshi and Thompson 1995).

At this point, we can only speculate about the reason(s) for why the evolution of increased competitive ability in the ACU and NCU populations occurred in a manner so different from that seen earlier in *D. melanogaster.* We believe there are three possible reasons for the observed discrepancy in correlated responses to selection for adaptation to larval crowding across the three species, and the three proposed explanations are not mutually exclusive. First, it is possible that the pattern of genetic variances and covariances among traits relevant to survival in a crowded culture is different from that in *D. melanogaster* in the two species studied here (*D. ananassae* and *D. n. nasuta*). For example, maybe the AB and NB populations do not harbour additive genetic variance for larval feeding rate. Such differences of genetic architecture could simply reflect between-species differences, being the result of historical selection pressures and contingencies. However, if that were the case, we would have expected the results from *D. ananassae* to be closer to *D. melanogaster,* as compared to *D. n. nasuta,* given the phylogenetic and ecological relationships of these species (see Introduction, last paragraph). A second, more likely, possibility is that these differences of genetic architecture between *D. ananassae* and *D. n. nasuta* on the one hand, and *D. melanogaster* on the other, are because the ACU and NCU populations represent selection on relatively recently wild-caught populations (see Materials and methods) whereas the *K*- and CU populations involved selection on populations that had already been in the laboratory for a large number of generations (Mueller and Ayala 1981; Joshi and Mueller 1996). There might be significant differences in the genetic architecture of traits related to fitness under larval crowding between wild and laboratory populations. Finally, it is possible that the discrepancies between the results of this study and those from earlier work on *D. melanogaster* are due to differences in the specific details of the respective maintenance regimes of selected populations in the various studies. Such small differences in selection regimes have been implicated in the different trajectories of pupation height in the *K-* versus the CU populations (Joshi et al. 2003). The *K*-populations were maintained on a serial transfer system in half-pint milk bottles with about 350 mL food per bottle (Mueller and Ayala 1981), while the CU populations were maintained in 6 dram vials (2.2 cm diameter) at a density of about 1000–1500 eggs per vial in about 5 mL of food. Unlike in our ACU and NCU populations, neither food amount nor number of eggs per vial were exactly controlled in CU maintenance. In the CU populations, eggs were laid by adult females onto thin films of food on four watch glasses per cage. These films of food with eggs on them were then sliced up and divided roughly equally among 40 vials that each contained about 3 mL of food (A. Joshi, *pers. obs.*). Thus, our ACU and NCU populations had a smaller absolute amount of food per rearing container (bottle/vial) than the *K*- and CU populations, and also differed from the latter in specific densities of eggs per mL of food. It is, therefore, possible that the time course of food depletion and nitrogenous waste build-up in the ACU and NCU cultures is somewhat different from that in the *K*- and CU populations. It has been shown theoretically that optimal feeding rates are likely to decline as the concentration of nitrogenous waste in the food increases (Mueller et al. 1995). Thus, at least in principle, it is possible that the optimal feeding rates in the ACU and NCU populations are actually less than they were for the *K*- and CU populations, and that is why increased feeding rates did not evolve in our experiments. Subsequent studies will attempt to discriminate between these various explanations. However, for the time being, we believe that these results underscore the need for a more nuanced understanding of adaptations to larval crowding in *Drosophila*, with a greater appreciation for the fact that increased competitive ability can be attained through the evolution of fairly different suites of traits. We also need to be cognizant of the fact that seemingly small differences of maintenance in otherwise similar selection regimes might mediate the evolution of very different trajectories in phenotypic space.

## Acknowledgements

We thank Larry Mueller for much helpful discussion, and N. Rajanna and M. Manjesh for help in the laboratory. A. Nagarajan thanks the Council of Scientific and Industrial Research, Government of India, for financial assistance in the form of Junior and Senior Research Fellowships. S. B. Natarajan was supported by a doctoral fellowship from the Jawaharlal Nehru Centre for Advanced Scientific Research. Shreyas Jois’ stay in the laboratory was supported by the joint Summer Research Fellowship programme of the three Indian science academies. This work was supported by funds from the Department of Science and Technology, Government of India, to A. Joshi. Preparation of the manuscript was supported in part by a J. C. Bose National Fellowship from the Department of Science and Technology, Government of India, to A. Joshi. The authors declare no competing interests.

